# Generative deep learning models for cognitive performance trajectories in real-world scenarios

**DOI:** 10.1101/2024.07.05.600608

**Authors:** Denis Expósito, Elina Maltseva, Carolina Sastre-Barrios, Iñigo Fernández de Piérola, Jesus M. Cortes

## Abstract

The increasing prevalence of cognitive disorders, such as Alzheimer’s disease, imposes significant challenges on healthcare systems and society. The ability to predict the future cognitive performance (CP) is crucial for professionals in neuropsychology, and real-world data emerges as an important source of complete and reliable information. However, its inherent complexities requires the use of advanced models to make predictions. To do so, we have implemented and compared three deep learning predictive strategies from CP trajectories: multilayer perceptron (MLP), convolutional neural networks (CNN) and long short-term memory (LSTM). The three models showed robustness on their predictions in different patient datasets. The CNN was the most suitable architecture due to its local pattern recognition capabilities and its robustness to overfitting. Therefore, professionals can have a complementary support for targeting treatment approaches to patients needs and anticipate undesired outcomes (e.g. cognitive impairment). Nonetheless, further studies are needed to validate whether neuropsychological interventions based on score predictions lead to improved intervention efficacy compared to traditional approaches for controlled patient groups.

## 1. Introduction

According to the Alzheimer’s Association, about 1 in 9 people aged 65 and older are living with Alzheimer’s disease (AD) or other cognitive disorders in the US. This proportion is expected to grow as the risk increases with advancing age, and more people will fall into the over-65 age group. These conditions not only affect individuals but also place a substantial burden on patient’s families, healthcare systems and society as a whole [1]. In 2022, caregivers alone offered an estimated 18 billion hours of informal (unpaid) assistance, which is valued at $339.5 billion, the equivalent of the Gross Domestic Product of a country like Colombia in the same year [2].

Cognitive performance (CP) encompasses various mental abilities, including memory, attention, and problem-solving skills. For individuals, particularly those being intervened by neuropsychologists, the trajectory of CP contains critical information of their overall well-being [3, 4]. Being able to anticipate changes in CP can have profound implications for patient care. Despite the significant importance of CP trajectories, their precise characterization over time at the individual level, in both healthy and pathological conditions, remains largely unknown.

CP assessment and the early prediction of cognitive impairment represent major challenges within neuropsychology. The development of predictive models has been mainly focused on biomarkers, clinical measurements such as neuroimaging, cognitive tests, and other parameters linked to the conversion from early cognitive impairment to dementia [5–10], even though there is still room for improvement.

It has been shown that traditional cohort studies have limitations, particularly in capturing the dynamic and diverse experiences of individuals in real-world settings [11–13]. Stringent inclusion and exclusion criteria, often imposed in controlled clinical environments, have a limited efficacy in generalizing to populations different from those recruited, and they may not fully represent the complexities of CP and cognitive impairment manifested in daily life. In this scenario, real-world data (RWD) can help to tackle this problem. The US Food and Drug Administration (FDA) defines RWD as “data relating to patient health status and/or the delivery of health care routinely collected from a variety of sources”. The rise in the adoption of digital sources of information like electronic health records, social media, and wearable devices has resulted in the generation and collection of massive quantities of RWD.

RWD allows the use of systems for early detection of cognitive decline and real-time recommendations for neuropsychologists to provide personalized interventions [14, 15], which have already shown positive results in therapy to delay the severity of dementia-related dependencies for institutionalization [16]. The digital transformation of information is also providing professionals with tools for data access and interventions from remote locations via cloud platforms, with increasing popularity [15, 17, 18]. However, RWD presents inherent difficulties due to its complexity, large-scale, and heterogeneous nature, and working with RWD is challenging [14]. Remarkable advances in the field of machine learning in the last decade, such as the implementation of deep neural network architectures, emerged as possible solutions to extract abstract representations and nonlinear relationships from complex data [14, 19].

In this study, we work with RWD from NeuronUP.com [20], an established platform providing hundreds of training materials (TMs) to work on cognitive stimulation and intervention. A NeuronUP Score [21] is calculated to keep track on patients’ CP over time. Recently, by using the NeuronUP score from TMs from different cognitive areas (domains), Camino-Pontes et al. [15] developed machine learning modeling to predict cognitive decline one year after being tested in NeuronUP patients. Here, we extend this study and develop deep learning (DL) models to predict future cognitive performance in patients using NeuronUP Scores. We first hypothesize that by developing and employing different DL architectures, we can construct a general model to accurately predict CP trajectories in different patients. An accurate model for prediction of cognitive decline holds great potential for the early detection of cognitive impairment conditions, enabling timely interventions and tailored approaches to meet the specific needs of individuals.

## 2. Materials and methods

Figure 1 shows a methodological sketch for this study, consisting of several steps to perform the data science analysis.

**Figure 1:**
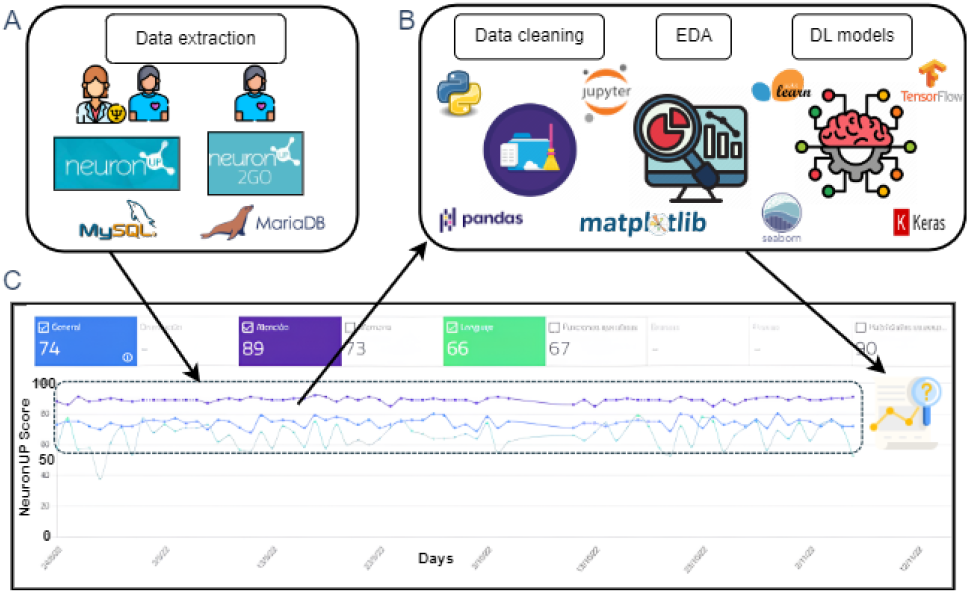
Data analysis and score prediction pipeline. **A**: Data was collected from NeuronUP using relational databases. **B**: A process of data cleaning followed by an exploratory data analysis (EDA) was done to the extracted raw data using the Jupyter notebook interface with Python language and some of its data analysis packages. Then a model development stage using ML and DL packages learned from the cleaned data and provided the predictions. **C**: Extracted data contained patient time series data on scores and cognitive function scores, which after the analysis future score values were predicted by the three models.

### 2.1. Ethical considerations

This study was exempted from an Ethics Review Panel as it makes use of low-risk retrospective research from existing collections of data that contain non-identifiable data, and where all participants provided information before collecting the data. In relation to the signed consent, all NeuronUP stimulation centres signed strict agreements for the use of data for research purposes. The agreement establishes that all responsible centres have, before collecting the data, to adequately inform on: (1) That the information collected will be used for various purposes related to research in prevention, diagnosis and treatment of diseases, and the promotion of health in society. (2) That there is the possibility of asking everything that the participant needs or does not understand.

(3) That participation is completely voluntary and that at any time the participant can withdraw without giving any explanation. In this case, the participant’s data will be removed from the NeuronUP platform. (4) That the NeuronUP data team will never have access to the participant’s personal data, and that their identity will be encrypted with an alphanumeric code and will not contain any personal or identifying data of the participant. (5) That the participant’s data will never be transferred to third parties or companies, although the possibility of NeuronUP collaborating with other partners in research projects is contemplated. All data used in this study come from participants who have given their consent by signing an electronic form.

### 2.2. NeuronUP Score

RWD from NeuronUP.com [20] was used. NeuronUP is a widely recognized neurorehabilitation and cognitive stimulation platform present in more than 30 different countries, and with more than 150K different users. NeuronUP.com offers a large repertoire of training materials (TMs) designed for cognitive stimulation and intervention, and generates a NeuronUP Score [21] to monitor patient performance longitudinally. This score is a novel quantitative index created to simplify a participant’s performance and to facilitate the comparisons in the follow-up, allowing longitudinal data of the same participant to be modelled while obtaining precise trajectories of individual performance.

The different NeuronUP TMs are grouped in 9 different cognitive domains hierarchically grouped including 29 subareas. First, each exercise for a given TM returns an exercise score (*s*_*j*_), which depends on the number of correct answers (*n*_0_), errors (*n*_1_), attempts (*n*_2_) and omissions (*n*_3_), being each of them weighted according to their importance in the specific TM (*p*_0_, *p*_1_, *p*_2_ and *p*_3_, respectively). After each exercise, we obtain:

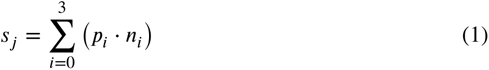

Which is multiplied by its difficulty (*d*_*i*_):

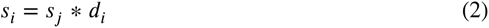

Then the TM score (*S*_*j*_) can be calculated as an average of all its exercise scores:

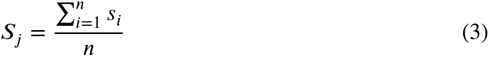

Within a TM, each cognitive function has a determined weight relative to its importance. Thus, it is possible to calculate a given score for a cognitive function (*S*_*l*_) from the score of all played TMs, and the weight of this specific cognitive function on these TMs (*w*_*i*_):

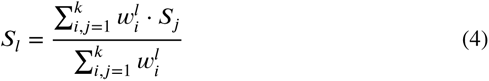

Finally, an overall score (S) can be obtained by combining each of the cognitive functions considering their impact based on its weights (%_*l*_):

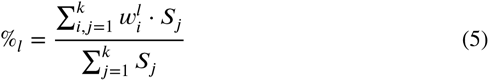

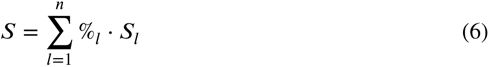

A score can be generated in two different situations: a therapy session with a professional using NeuronUP or the patient exercising alone at home using NeuronUP 2GO.

### 2.3. Data engineering

NeuronUP data was collected using MySQL and MariaDB relational database management systems and the SQL language for querying. Raw data was divided into one file for 2014-2020 period and another for 2021-2023 period, as data size was too large for a unique file. The remaining parts of the project were executed using Python programming language in the Jupyter notebook development environment. For reproducibility, the notebooks used in this study are available in the GitHub repository (the repository can be accessed at: https://github.com/deexposito/NeuronUP_Score_Prediction). Due to confidentiality reasons, the underlying data cannot be shared publicly, but detailed descriptions of the data and preprocessing steps are provided within the article. After merging the two data subsets, we performed a quick revision of their attributes, with a total of *N* = 19642 patients.

### 2.4. Data preprocessing

Major steps for data preprocessing were:

#### Missing values and outliers consideration

As gender and birth-date variables were considered as basic information for the model, records with missing values for one of them were removed. Values from score and area score outside the [0, 1] range were considered as outliers and excluded from the study. This preprocessing step ensured that the dataset used for analysis was clean and reliable, minimizing the risk of introducing bias or inaccuracies due to incomplete or erroneous data.

#### Data formatting

A time series model requires to have unique rows per time unit (per day and patient in this case, as we consider each patient as isolated data). However, the area ID variable forced the dataset to have more than one row per patient and day combination (one for each area), so we established each area as a new attribute with area score values, and merged all area scores in the same day, and dropped the area ID and area score columns. Day values were not consecutive due to the intrinsic irregularity of cognitive stimulation (i.e. patients cannot be forced to play every day), so we compacted the days into weeks using mean score values for each week. As the model considers each patient as a separated dataset, we needed patients with a minimum of weeks played, and we filtered them using patients with minimum the median value (7 weeks) and reducing the number of patients to 10113.

#### Binning categories into groups

We considered the age group as an interesting variable for modeling, so we grouped the birth-date attribute into a new one ‘age group’ with five categories: pre-scholars (0-5 y/o), children (6-11 y/o), teenagers (12-18 y/o), young adults (19-34 y/o), middle-aged adults (35-64 y/o) and senior adults (>65 y/o).

#### Emerging missing values

When we structured the data into unique patient-day combinations, new missing values emerged in each of the cognitive areas, as they were not present in every time point. Percentages of missing values in each cognitive function are: executive functions (11,18%), attention (12,19%), memory (39,08%), language (43,5%), gnosis (59,59%), visuospatial skills (60,19%), social cognition (90,38%), orientation (90,7%) and praxis (93,27%). KNN imputation was done with 3 neighbors (k), allowing to rescue 6 of the 9 cognitive functions to be used for subsequent analysis: attention, executive functions, gnosis, language, memory and visuospatial skills. The number of patients was 7253. We conducted empirical testing to determine the optimal k value using different k values (1, 3 and 5). For each k, we examined the correlations between imputed variables and other variables in the dataset to ensure that the imputed values preserved the relationships in the data.

### 2.5. Exploratory data analysis

After cleaning the dataset, we performed an exploratory data analysis (EDA, see Figure S1 for detailed visualizations) to gain a deeper understanding of the data. Before EDA, we checked again the summary statistics of our data and ensured the presence of 10113 patients, 2 genders, and the general score and all area scores between 0 and 1.

#### Gender and age group distributions

Genders were distributed as 56% male (n=5660) and 44% female (n=4453). Regarding age groups, we had 2.7% preescolars (n=273), 10.6% children (n=1056), 16.5% teenagers (n=1639), 13.9% young adults (n=1384), 28.4% middle-aged adults (n=2829) and 27.9% senior adults (n=2772).

#### Score and area scores distributions

For all the general score and the area scores, each total of the 8 Saphiro-Wilk normality tests were significantly non-Gaussian (p-value < 0.01). No correlation was found between score and gender or with age group.

### 2.6. Time series modeling

#### Cohort selection

To evaluate reproducibility in the model, we performed a cohort selection in 4 groups with stratification for gender and age group, so all groups have an equal proportion of them. One split was done on the initial dataset and a Chi-square test showed no significant differences between groups in gender and age group distributions. Another split was done in each of the two subsets and the same test in both cases again with no significant differences.

#### Three different deep-learning models for time-series prediction

One cohort (cohort 1) was used for model training and evaluation to optimize the hyper-parameters. The remaining three cohorts (cohorts 2, 3 and 4) were used for evaluation reproducibility of the model. Three different neural network approaches were conducted in parallel using the Keras Python library: multilayer perceptron (MLP), long short-term memory (LSTM) and convolutional neural network (CNN) (their neural network architectures can be found in Figure 2).

**Figure 2:**
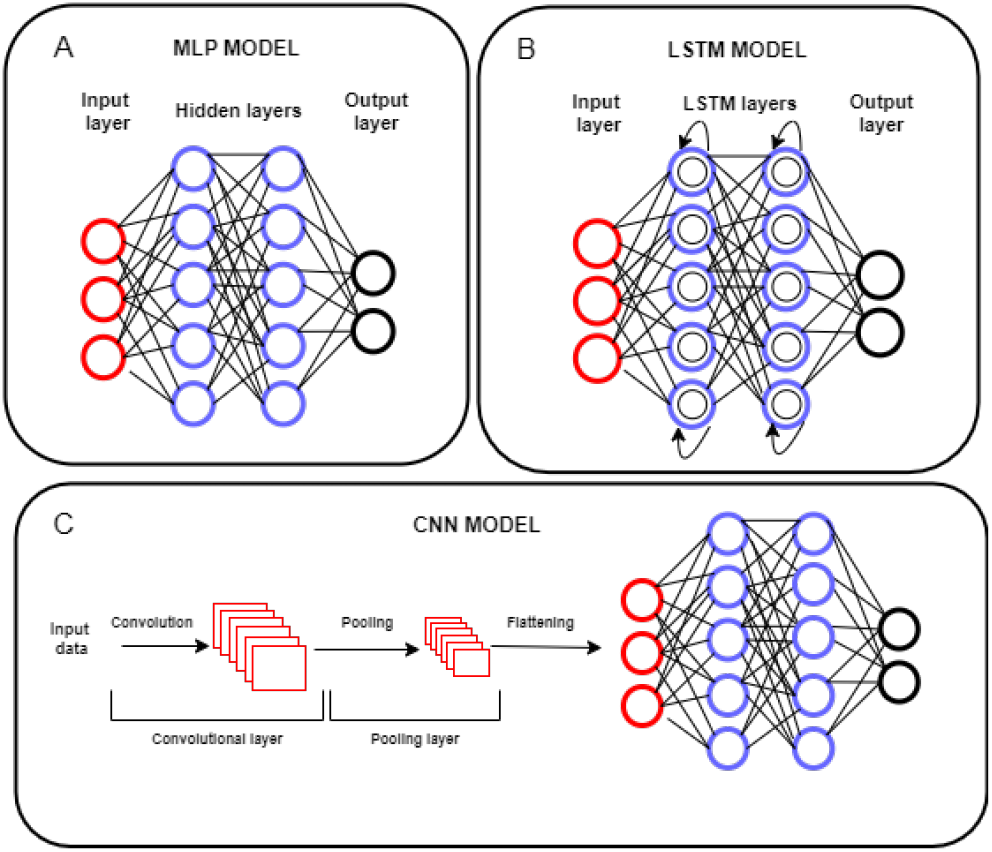
Neural network architectures, in three different models. **A**: MLP, sketched by an input layer, hidden layers and an output layer. **B**: The LSTM model achieves a short-term memory characteristic of recurrent neural network plus a long-term memory solving their vanishing gradient problem. **C**: The CNN architecture requires a process of convolution and pooling through a convolutional and pooling layer, respectively, and then the output is flatten into a MLP structure.

<u>MLPs</u> are feedforward neural networks consisting of several layers of interconnected nodes or neurons, where information flows in one direction from an input layer, through one or more hidden layers, into an output layer. Computations in every neuron depend on a series of parameters: an activation function *s*, a weight between pairs of neurons *w*, a neuron’s bias vector *b*, and the activation of the previous input *x* or hidden *h* layer [22].

<u>LSTMs</u> are a variation of recurrent neural networks, which is limited to memorize short-term past information due to a vanishing gradient problem, while this variation allows to overcome this problem and recall long-term time series data. In LSTM computations, each neuron has several parameters: an input gate *i*, a forget gate *f*, an output gate *o*, a cell state *C*, and a hidden state *h* [23].

<u>CNNs</u> assume that input sequences have image structures. Multiple filters extract features from data as feature maps (convolution), where a pooling operation performs dimensionality reduction, and then these pooled feature maps are flattened into a one-dimensional vector to be fed to a neural network. In CNN computations, each neuron in a convolutional layer has parameters: a filter (or kernel) *K*, a bias *b*, and an activation function *s* [23].

#### Reshaping time series data

For every patient, data was transformed into a 2D supervised learning format using a window step size of two weeks, so each prediction takes into account the two previous weeks. LSTM and CNN models needed a 3D structure [samples, time steps, features], while MLP model required to flatten the input samples [samples, features].

#### Hyper-parameter tuning

To determine the optimal model hyper-parameters we performed a time series cross-validation and a grid search in the first cohort to train each model with all possible hyperparameter combinations using 75% of the dataset for training and the remaining 25% for testing.

The selection criteria for the hyper-parameters were based on the following considerations:

##### 1) Performance Metric

We used a custom loss function using the root mean squared error (RMSE) while incorporating a penalty for predictions falling outside the natural [0, 1] range of the score values. This ensures that the model not only fits the data well but also respects the valid range of the predictions.

##### 2) Time Series Cross-Validation

To account for the temporal dependencies in our dataset, we employed time series cross-validation. This method helps in providing a more realistic estimate of the model’s performance on unseen data, as it preserves the temporal order and avoids data leakage.

##### 3) Grid Search

We performed an exhaustive grid search over a defined set of hyper-parameter values. This systematic approach allowed us to evaluate the performance of each possible combination of hyper-parameters and identify the optimal settings.

##### 4) Loss Plots

During the evaluation phase, we generated loss plots to visualize the performance of different hyperparameter values. These plots identify not only the hyperparameter values that minimized the loss but also those that provided stable performance with fewer outliers.

By combining these methods and criteria, we ensured that the selected hyper-parameters were both optimal in terms of performance and robust in their predictive capabilities.

#### Prediction accuracy

For each model with their optimal hyper-parameters predictions were made at the patient-level for all other three cohorts. A short-sentence of 20 previous time points (values of score) were used for training, aiming to predict the following 10 points and compare them to the true values. To evaluate prediction, we calculated RMSE that was used as the main model performance metric, although other metrics were also used such as the forecast bias (which measures the systematic deviation from true values), the forecast interval coverage (assessing the proportion of true values falling within the prediction ±0.1), and the prediction direction accuracy (measuring the proportion of directional (increase or decrease) predictions of future values [24]). We analyzed the performance differences between two different strategies: intra-model, comparing all metrics (except PDA) for the same model in the three different datasets to evaluate their reproducibility, and inter-model, comparing metrics between models to discover the best modeling approach. A Kruskal-Wallis statistic was used for testing the intra-model scenario, while a Friedman test was used in the inter-model scenario (where we merged the data from the three datasets), and a Nemenyi post-hoc pairwise comparison was considered to assess significant comparisons.

### Statistical analyses

To test for differences in performance between intramodel data, inter-model data, and to check for Gaussianity in the distribution of values for each model’s performance metric, we used respectively a Kruskal-Wallis test, a Friedman test, and a Saphiro-Wilk test, all of them implemented in the *SciPy*.*stats* library (version 1.11.4) and available at https://scipy.org/. The pairwise post-hoc analysis was done using the Nemenyi test implemented in the *Scikit-posthocs* package (version 0.9.0) and available at https://pypi.org/project/scikit-posthocs/. Differences between gender scores were tested using the Wilcoxon signed-rank test, and for age groups we used again the Kruskal-Wallis test and then a Dunn’s test using *SciPy*.*stats* and *statsmodels* module (version 0.13.5). All these analysis were performed in Python 3.11.5 available at https://www.python.org/.

## 3. Results

The ReLU activation function combined with a learning rate of 0.001 yielded the best performance (see Table 1). Variations in the number of layers did not significantly affect loss values, leading us to adopt the minimum complexity case to optimize computational efficiency during training.

**Table 1.** Hyper-Parameter (HP) tuning of models.

| HP Name | HP Range | MLP | LSTM | CNN |
| --- | --- | --- | --- | --- |
| Activation function | [ReLU, SeLU, Tanh] | ReLU | ReLU | ReLU |
| Dropout rate | [0, 0.05, 0.1] | 0.05 | 0.05 | 0.05 |
| Learning rate | [0.001, 0.01, 0.1] | 0.001 | 0.001 | 0.001 |
| N<sup>o</sup> dense layers | [1, 2, 3] | 1 | 1 | 1 |
| N<sup>o</sup> LSTM layers | [1, 2, 3] | – | 1 | – |
| N<sup>o</sup> Conv1D layers | [1, 2, 3] | – | – | 1 |

No significant differences were found in any of the three models for any of the evaluation metrics by considering the three evaluation cohorts, which highlights that the prediction accuracy was model-independent, emphasizing the reproducibility of our DL models for the different scenarios studied (see Tables 2 and 3).

**Table 2.**
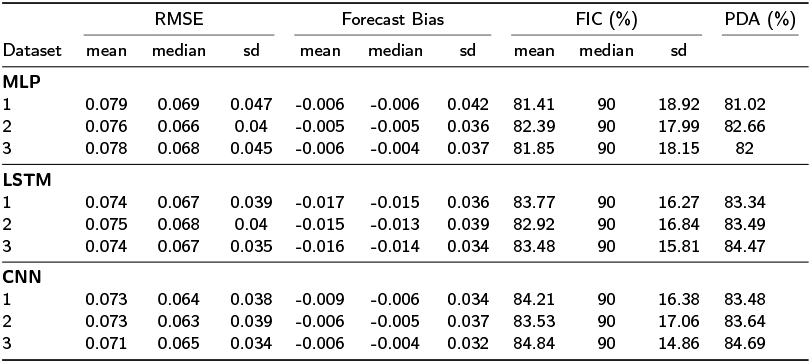
Performance metrics for MLP, LSTM, and CNN models. Three evaluation cohorts were used (for each cohort, n = 1803)

| Dataset | RMSE |  |  | Forecast Bias |  |  | FIC (%) |  |  | PDA (%) |
| --- | --- | --- | --- | --- | --- | --- | --- | --- | --- | --- |
|  | mean | median | sd | mean | median | sd | mean | median | sd |  |
| **MLP** |  |  |  |  |  |  |  |  |  |  |
| 1 | 0.079 | 0.069 | 0.047 | -0.006 | -0.006 | 0.042 | 81.41 | 90 | 18.92 | 81.02 |
| 2 | 0.076 | 0.066 | 0.04 | -0.005 | -0.005 | 0.036 | 82.39 | 90 | 17.99 | 82.66 |
| 3 | 0.078 | 0.068 | 0.045 | -0.006 | -0.004 | 0.037 | 81.85 | 90 | 18.15 | 82 |
| **LSTM** |  |  |  |  |  |  |  |  |  |  |
| 1 | 0.074 | 0.067 | 0.039 | -0.017 | -0.015 | 0.036 | 83.77 | 90 | 16.27 | 83.34 |
| 2 | 0.075 | 0.068 | 0.04 | -0.015 | -0.013 | 0.039 | 82.92 | 90 | 16.84 | 83.49 |
| 3 | 0.074 | 0.067 | 0.035 | -0.016 | -0.014 | 0.034 | 83.48 | 90 | 15.81 | 84.47 |
| **CNN** |  |  |  |  |  |  |  |  |  |  |
| 1 | 0.073 | 0.064 | 0.038 | -0.009 | -0.006 | 0.034 | 84.21 | 90 | 16.38 | 83.48 |
| 2 | 0.073 | 0.063 | 0.039 | -0.006 | -0.005 | 0.037 | 83.53 | 90 | 17.06 | 83.64 |
| 3 | 0.071 | 0.065 | 0.034 | -0.006 | -0.004 | 0.032 | 84.84 | 90 | 14.86 | 84.69 |

**Table 3.** Kruskal-Wallis test to evaluate model reproducibility. Each intra-model data consisted of the three evaluation cohorts concatenated (n = 1813×3).

| Intra-model data | Kruskal-Wallis test |  |  |  |  |  |
| --- | --- | --- | --- | --- | --- | --- |
|  | RMSE |  | Forecast Bias |  | FIC |  |
|  | Statistic | p-value | Statistic | p-value | Statistic | p-value |
| MLP | 0.583 | 0.75 | 1.335 | 0.51 | 0.798 | 0.67 |
| LSTM | 0.041 | 0.98 | 0.826 | 0.66 | 1.035 | 0.60 |
| CNN | 0.136 | 0.93 | 2.377 | 0.30 | 0.697 | 0.71 |

For model comparisons, we first tested whether our data was significantly differing from Gaussianity, as most of the classical statistical tests assume that data follows a Gaussian distribution. Our results show that the hypothesis of a Gaussian distribution in prediction values was rejected in all models for all metrics as shown in Table 4. Significant differences were found between models in the Friedman test for RMSE, forecast bias and FIC, and the post-hoc analysis focused these differences for lower RMSE in the CNN model, higher absolute forecast bias in the LSTM model and divergence in FIC values for all models (MLP<LSTM<CNN) (see Tables 5 and 6). A visual comparison of predictions using every model in every gender-age group possible combination is shown in Figure 3.

**Table 4.** Saphiro-Wilk test for normality of models’ evaluation metrics. Each model data consisted of the three evaluation cohorts concatenated (n = 1813×3).

| Model | Saphiro-Wilk test |  |  |  |  |  |
| --- | --- | --- | --- | --- | --- | --- |
|  | RMSE |  | Forecast Bias |  | FIC |  |
|  | Statistic | p-value | Statistic | p-value | Statistic | p-value |
| MLP | 0.835 | <0.001 | 0.937 | <0.001 | 0.853 | <0.001 |
| LSTM | 0.871 | <0.001 | 0.928 | <0.001 | 0.86 | <0.001 |
| CNN | 0.887 | <0.001 | 0.953 | <0.001 | 0.854 | <0.001 |

**Table 5.**
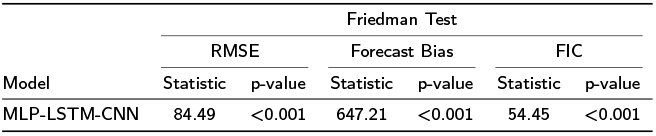
Friedman Test for differences between models in terms of evaluation metrics. The comparison was done considering each model dataset as the three concatenated cohorts (n = 1813×3).

**Table 6.** Nemenyi post-hoc analysis for pairwise differences in evaluation metrics between models. Each comparison was done considering each model dataset as the three concatenated cohorts (n = 1813×3).

| Pairwise comparisons | Nemenyi post-hoc analysis. |  |  |
| --- | --- | --- | --- |
|  | RMSE | Forecast Bias | FIC |
|  | p-value | p-value | p-value |
| MLP-LSTM | 0.687 | 0.01 | 0.011 |
| MLP-CNN | 0.001 | 0.9 | 0.01 |
| LSTM-CNN | 0.001 | 0.01 | 0.046 |

**Table 7.** Dunn’s Post-Hoc Test Results: Comparison of scores between different age groups. We can observe significant differences in all age group combinations except between Children and Young Adults.

| Group Comparison | Dunn's Post-Hoc Test |  |
| --- | --- | --- |
|  | p-value | p-adjusted |
| Prescolar(0-5) vs Children(6-11) | <0.001 | <0.001 |
| Prescolar(0-5) vs Teenagers(12-18) | <0.001 | <0.001 |
| Prescolar(0-5) vs Young Adults(19-34) | <0.001 | <0.001 |
| Prescolar(0-5) vs Middle-Aged Adults(35-64) | <0.001 | <0.001 |
| Prescolar(0-5) vs Senior Adults(>65) | <0.001 | <0.001 |
| Children(6-11) vs Teenagers(12-18) | <0.001 | <0.001 |
| Children(6-11) vs Young Adults(19-34) | 0.415 | 1.000 |
| Children(6-11) vs Middle-Aged Adults(35-64) | <0.001 | <0.001 |
| Children(6-11) vs Senior Adults(>65) | <0.001 | <0.001 |
| Teenagers(12-18) vs Young Adults(19-34) | <0.001 | <0.001 |
| Teenagers(12-18) vs Middle-Aged Adults(35-64) | <0.001 | <0.001 |
| Teenagers(12-18) vs Senior Adults(>65) | <0.001 | <0.001 |
| Young Adults(19-34) vs Middle-Aged Adults(35-64) | <0.001 | <0.001 |
| Young Adults(19-34) vs Senior Adults(>65) | <0.001 | <0.001 |
| Middle-Aged Adults(35-64) vs Senior Adults(>65) | <0.001 | <0.001 |

**Figure 3:**
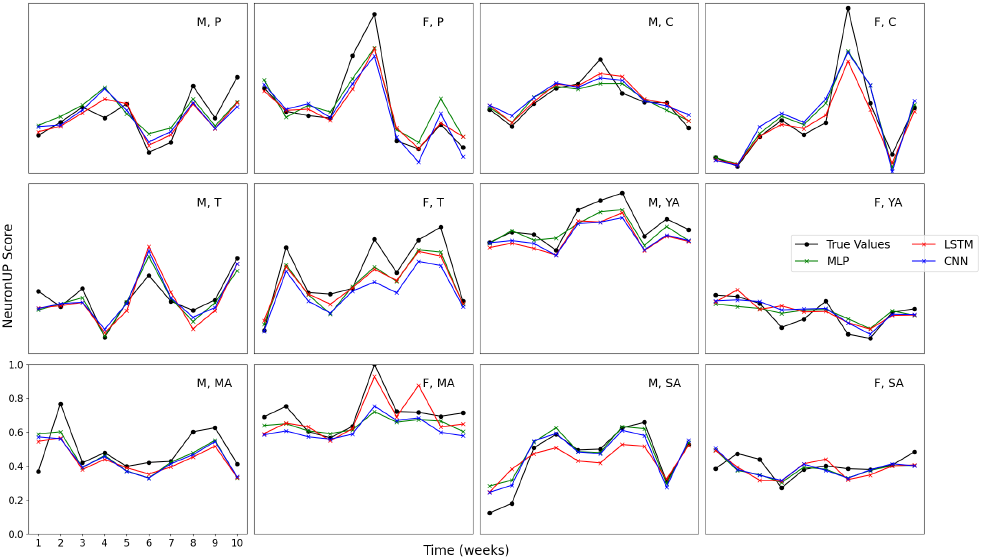
Model predictions in all types of patients for gender and age group against true values. For all panels, training was performed over 20 time points and testing was performed over 10 time points (used for predictions). Here, we depict a single trail prediction for illustration purposes. Notice that for each of the panels, the score was calculated over a different intervention. M: Male, F: Female, P: Pre-scholar, C: Children, T: Teenager, YA: Young Adult, MA: Middle-Aged Adult, SA: Senior Adult.

We also analyzed the differences in scores between genders and age groups using non-parametric tests, and significant differences were revealed in genders (Wilcoxon ranksum statistic = 2.2 × 10^10^, p-value = *<* 0.001). The same scenario of differences is produced between age groups (Kruskal-Wallis statistic = 1199.76, p-value = *<* 0.001 ;for the posthoc analysis, see 7).

## Discussion

Three different deep learning model architectures were used for prediction of cognitive performance trajectories, namely, multilayer perceptron (MLP), convolutional neural networks (CNN) and long short-term memory (LSTM). We have assessed their reproducibility and predictive performance by using three different evaluation metrics: RMSE, forecast bias and forecast interval coverage. All three models needed to be adjusted from the neural network architecture, as the number of hidden layers, neurons in each layer, the activation function, dropout rate and learning rate, providing flexibility to adapt to different datasets. The models learn to extract non-linear relationships between input and predictive variables, thus capturing temporal patterns and make accurate predictions.

MLPs use layers of fully connected neurons, and stand out for their simplicity, even though they lack temporal memory. LSTMs, as recurrent neural networks, capture temporal patterns, and include memory units with capacity for long-term information, which on the other hand may require more training time. CNNs have convolutional layers that scan the input data to capture complex patterns and relationships, are computationally efficient, and well-suited for short-term information, while they struggle with long-term dependencies.

Our results show a consistent pattern across all three models, indicating that the chosen hyper-parameter values were robust, and supporting the credibility of their predictive capability. Hyper-parameter choices were made with consideration of performance, prevention of over-fitting, and computational efficiency.

For each model, our intra-model comparisons revealed no significant differences in any of the evaluation metrics across three different evaluation cohorts. The consistency in model performance across multiple datasets supports the robustness and reproducibility of our DL models. Our finding remark the reliability of our models’ predictions and underscores their suitability for different neuropsychological clinical contexts in the platform.

In the inter-model comparison, the CNN model outperformed the MLP and LSTM models in RMSE and FIC evaluation metrics. The reason for the CNN model’s better performance may lie in its architecture:

1. Local pattern recognition: CNNs are particularly effective at identifying local patterns thanks to their convolutional layers, which apply filters to the input data to detect trends and shifts over time.
2. Robustness to overfitting: CNNs’ pooling layers reduce the spatial dimensions of the data while preserving important features, which helps to mitigate the risk of overfitting.

The superior performance of the CNN model suggests that the temporal dynamics of cognitive performance trajectories captured in our dataset are likely driven by short-term changes.

Moreover, the LSTM model, despite its capability to capture long-term dependencies, showed higher absolute forecast bias compared to the other models. A possible explanation may arise from the inherent characteristics of LSTM networks, such as their ability to capture long-term dependencies in sequential data, as it could potentially lead to over-fitting or biases in certain scenarios, resulting in higher forecast bias. Based on these observations, we considered the CNN model as the most suitable option for the data obtained from the NeuronUP platform.

A comparison of all different predictions across different gender-age group combinations provides another interesting result from our study, in particular, that the three models accurately predict future score values. In consequence, clinical professionals can better anticipate to future cognitive decline, having time in advance to evaluate their intervention strategies. In addition, our analysis revealed significant differences in cognitive scores between genders and different age groups. These gender and age group differences underscore the importance of considering demographic variables in the analysis and interpretation of cognitive performance data.

However, the results presented in this study are specific to the NeuronUP platform data. When considering the generalizability of these findings to other datasets and real-world settings, several factors must be taken into account:

- Data characteristics: This NeuronUP dataset has unique characteristics in terms of cognitive performance measures and patient demographics. The performance of the model may vary when applied to different datasets.
- Model adaptation: The model may require new training and hyper-parameter tuning processes when applied to new datasets, and performance may vary depending on the flexibility of the model.
- Scalability: Empirical validation with large-scale data will be necessary to prove if its computational efficiency is scalable to larger datasets.

Our study also face some limitations. First of all, since scores are not available for all TMs and all subjects, it is not feasible to apply well-known techniques, such as Principal Component Analysis, to the original data to obtain a dimensionality reduction across different TMs. Second, RWD is also heterogeneous in the number of TMs within each cognitive domain, and future studies are needed for assessing the internal consistency of each domain. Third, there exist missing values in some of the co-variables such as the patients’ educational level, making not feasible to control our results by this co-variable.

Overall, we have shown that generative deep learning models produce accurate predictions of NeuronUP scores, which opens promising avenues for decision support to neuropsychologists in monitoring patients’ cognitive performance. Further studies should validate whether neuropsychological interventions based on score predictions, compared to interventions without score prediction, result in improved intervention efficacy for controlled patient groups. The superior performance of the CNN model highlights the importance of selecting appropriate model architectures and hyperparameters according to the characteristics of the input data.

## CRediT authorship contribution statement

**Denis Expósito:** Designed research, Performed the analyses, Made the figures, Drafted the first manuscript, Wrote the manuscript. **Elina Maltseva:** Data engineering, Data curation, Wrote the manuscript. **Carolina Sastre-Barrios:** Neuropsychologist, Wrote the manuscript. **Iñigo Fernández de Piérola:** Neuropsychologist, Wrote the manuscript. **Jesus M. Cortes:** Designed research, Supervised the research, Wrote the manuscript.

## Acknowledgements

D.E. acknowledges financial support from an internship grant from NeuronUP. This research has been funded by NeuronUP.com from its own resources, and from Centro para el Desarrollo Tecnológico Industrial de España (CDTI) (grant no. IDI-20210936).

---

**Figure S1:**
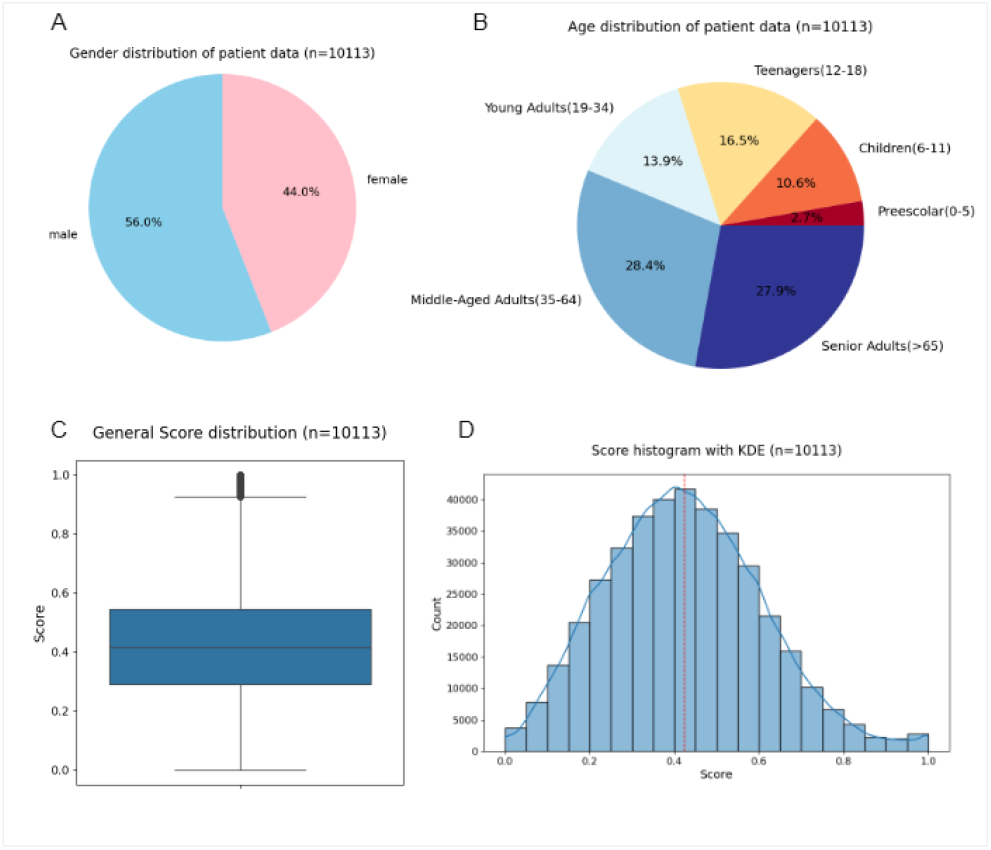
Demographic information and different statistics on cognitive performance. **A**: Gender distribution. **B**: Age-group distribution. **C**: NeuronUP Score box plot across time points and different subjects. **D**: NeuronUP Score histogram across time points and different subjects..

