## Supplementary figures and images for "Generative deep learning models for cognitive performance trajectories in real-world scenarios"

### Suplemmentary Figure 1

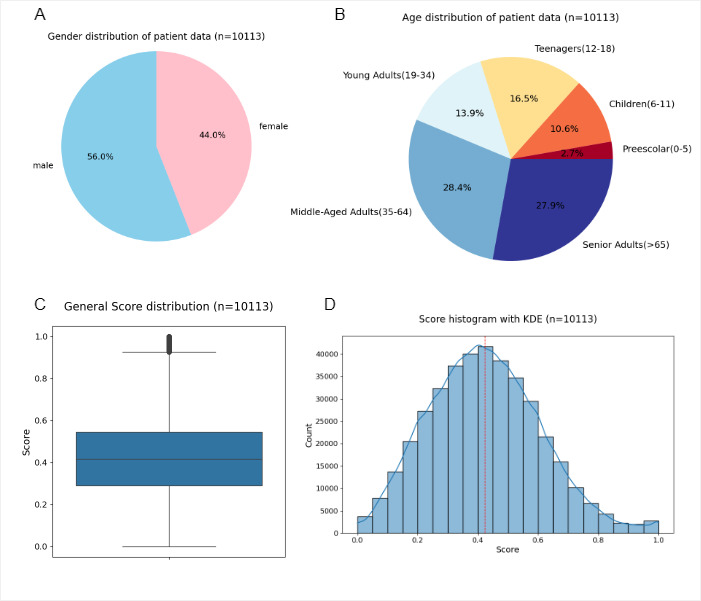
